# Sex differences in avoidance behavior and cued threat memory dynamics in mice: Interactions between estrous cycle and genetic background

**DOI:** 10.1101/2023.09.23.559127

**Authors:** Garret L. Ryherd, Averie L. Bunce, Haley A. Edwards, Nina E. Baumgartner, Elizabeth K. Lucas

## Abstract

Anxiety disorders are the most prevalent mental illnesses worldwide, exhibit high heritability, and affect twice as many women as men. To evaluate potential interactions between genetic background and cycling ovarian hormones on sex differences in susceptibility to negative valence behaviors relevant to anxiety disorders, we assayed avoidance behavior and cued threat memory dynamics in gonadally-intact adult male and female mice across four common inbred mouse strains: C57Bl/6J, 129S1/SVlmJ, DBA/2J, and BALB/cJ. Independent of sex, C57Bl/6J mice exhibited low avoidance but high threat memory, 129S1/SvlmJ mice high avoidance and high threat memory, DBA/2J mice low avoidance and low threat memory, and BALB/cJ mice high avoidance but low threat memory. Within-strain comparisons revealed reduced avoidance behavior in the high hormone phase of the estrous cycle (proestrus) compared to all other estrous phases in all strains except DBA/2J, which did not exhibit cycle-dependent behavioral fluctuations. Robust and opposing sex differences in threat conditioning and extinction training were found in the C57Bl/6J and 129S1/SvlmJ lines, whereas no sex differences were observed in the DBA/2J or BALB/cJ lines. C57Bl/6J males exhibited enhanced acute threat memory, whereas 129S1/SvlmJ females exhibited enhanced sustained threat memory, compared to their sex-matched littermates. These effects were not mediated by estrous cycle stage or sex differences in active versus passive defensive behavioral responses. Our data demonstrate that core features of behavioral endophenotypes relevant to anxiety disorders, such as avoidance and threat memory, are genetically driven yet dissociable and can be influenced further by cycling ovarian hormones.

## INTRODUCTION

Anxiety disorders, such as generalized anxiety disorder, panic disorder, and phobia, are the leading cause of mental illness, with a lifetime prevalence rate of 16% worldwide and 31% in the United States alone (Kessler et al., 2009; Kessler et al., 2005). Women are twice as likely as men to experience an anxiety disorder (Kessler et al., 2005; Kessler et al., 2006; Kessler et al., 1995), and epidemiological data further indicate that this increased susceptibility primarily occurs during years of reproductive viability (Hantsoo and Epperson, 2017). Greater symptom severity also leads to poorer quality of life in women with these conditions (Altemus et al., 2014; Maeng and Milad, 2015), an outcome driven by regulation of symptoms across the menstrual cycle in some individuals (Green and Graham, 2022). Interestingly, heritability estimates from both twin and genome-wide association studies indicate increased heritability of anxiety disorders in women compared to men (Duncan et al., 2018; Taylor et al., 2008), as well as strong associations between genetic diversity in sex hormone signaling and anxiety disorder risk (Levey et al., 2020). This clinical reality suggests that cycling ovarian hormones may interact with genetic composition to promote susceptibility and resilience to anxiety disorders.

While anxiety is a complex internal state that remains poorly understood, anxiety disorders are characterized by exaggerated emotional responses to real or perceived threats (Barroca et al., 2022). As such, key endophenotypes of anxiety disorders can be examined in preclinical mouse models, which are a powerful tool for disentangling the biological mechanisms governing behavioral traits. Traits relevant to anxiety disorders fall under the negative valence systems research domain criteria and include behavioral responses to acute threat, potential threat, sustained threat, loss, and frustrative nonreward (Cuthbert and Insel, 2013). Like humans, genetically divergent inbred mouse lines exhibit a spectrum of behavioral phenotypes, including behaviors relevant to negative valence systems and responses to anxiolytic compounds (Crawley et al., 1997). While sex differences in such behaviors have been found to diverge across mouse genetic backgrounds (Archer, 1977; Bolivar et al., 2001; Eltokhi et al., 2020), the literature paints a highly inconsistent picture regarding the impacts of sex and ovarian hormone levels in rodents (Bauer, 2023; Kokras and Dalla, 2014). However, as very few strain comparisons of negative valence behaviors have considered the interaction of cycling ovarian hormones and genetic composition, we hypothesized that inconsistencies in the literature may be due to a lack of consideration of mouse strain.

To address this gap in knowledge, behavioral responses across three categories of negative valence systems were measured in four common inbred mouse strains: C57Bl/6J, 129S1/SVlmJ, DBA/2J, and BALB/cJ. The rodent estrous cycle, which mirrors the reproductive hormone fluctuations of the human menstrual cycle (Knobil and Neill, 1994), was tracked throughout behavioral testing to stratify data to reproductive cycle phase. Responses to potential threat were assayed with center avoidance in the open field test and open arm avoidance in the elevated plus maze, while responses to acute and sustained threat were assayed with conditioned responses to auditory threat conditioning before and after extinction training. We demonstrate striking strain differences in avoidance and conditioned threat, with sex differences and estrous cycle regulation of these negative valence behaviors varying across mouse strains.

## MATERIALS AND METHODS

### Animals

C57Bl/6J (JAX #000664), 129S1/SVImJ (JAX #002448), DBA/2J (JAX #000671), and BALB/cJ (JAX #000651) breeding pairs were ordered from the Jackson Laboratory (Bar Harbor, ME, USA). Gonadally-intact male and female progeny aged 4-5 months were used for all experiments. Mice were housed 2-4 per cage with corncob bedding and *ad libitum* access to standard chow (LabDiet Rodent 5001) and water in an environmentally controlled husbandry room maintained on a 12h:12h light:dark cycle (lights on at 08:00 AM). To limit non-specific effects of litter on adult behavior (Valiquette et al., 2023), each group was comprised of ≥ 5 litters and, on average, 3 mice per litter. All experiments were conducted in accordance with the National Institutes of Health *Guide for the Care and Use of Laboratory Animals* and approved in advance by the Institutional Animal Care and Use Committee of North Carolina State University.

### Estrous Cycle Monitoring

Estrous cycle was monitored throughout the duration of experimental procedures through categorization of vaginal cytology (Cora et al., 2015). Cells were collected daily between 10:00-10:30 AM with vaginal lavage of 50 µl saline, and male animals were mock lavaged (anogenital area touched with pipette tip) to account for the additional handling received by females. Vaginal cytology was visualized with 1% toluidine blue staining. Estrous cycle stage was categorized as follows: proestrus, nucleated epithelial cells; estrus, cornified cells; diestrus, presence of leukocytes.

### Behavioral Testing – General Considerations

All behavioral testing occurred between 12:00-07:00 PM. All mice were handled and tested by the same researcher (G.L.R.). To minimize effects of sex pheromones on behavioral assays, male and female mice were tested in separate cohorts, and equipment was broken down and thoroughly cleaned between cohorts. Strains were interleaved among cohorts for a total of 7 male cohorts and 8 female cohorts, and the behavioral testing sequence alternated between male and female cohorts. Mice were habituated to an airlock outside the behavioral testing suite 30 minutes prior to each behavioral test. All mice went through an identical sequence: Days 1-3, handling; Day 4, open field test; Day 5, elevated plus maze; Day 6 auditory threat conditioning; Day 7, extinction training; Day 8, extinction training; Day 15, spontaneous recovery, renewal, and reminder shocks; Day 16, reinstatement. Due to a change in lighting conditions for the open field test and elevated plus maze, the first 26 mice were dropped from tracking analysis but included in analysis of threat memory dynamics. All experiments were analyzed blind to sex, but discrete coat colors precluded blinding to strain.

### Open Field Test

The open field test was conducted in an opaque gray arena (45 x 45 x 40 cm; Panlab, Harvard Apparatus, Holliston, MA) over a 30 min session. The arena was indirectly lit at 85 lux. Mice were placed in the corner of the arena and recorded with an overhead camera. Tracking was analyzed offline with AnyMaze software (Stoelting, Wood Dale, IL), and the center zone was set as the inner 225 cm^2^ of the arena.

### Elevated Plus Maze

The elevated plus maze consisted of two open arms (15 cm), two black opaque enclosed arms (15 cm), and a gray opaque floor (65 x 65 x 55 cm; Panlab). The arena was indirectly lit with the center and open arms at 120 lux. Mice were placed in the center of the maze facing the open arms and recorded with an overhead camera for 5 min. Tracking was analyzed offline with AnyMaze software.

### Cued Threat Memory Dynamics

Cued threat conditioning, extinction training, spontaneous recovery, renewal, and reinstatement were conducted in a near infrared video Habitest modular operant chamber housed within a sound-attenuating cubicle (Coulbourn, Holliston, MA). The conditioning context consisted of two clear Plexiglas walls and two stainless steel walls with an aluminum shock grid floor, and 70% ethanol was used as the odorant. The conditioned stimulus (CS) was an auditory tone (20 s, 2 kHz, 65 dB), and the unconditioned stimulus (US) was a footshock (0.5 mA, 2 s). For threat conditioning, mice received six co-terminating CS-US pairings (100 s inter-trial interval) after a 240 s habituation period. For extinction training, the walls and floor of the chamber were covered with white inserts adorned with red polka dots, and isopropanol was used as the odorant. Massed extinction was conducted over a two-day period. On each day, mice received 20 unreinforced presentations of the CS (60 s inter-trial interval) after a 180 s habituation period. The first CS block of extinction day one was used to assay acute threat memory recall. Seven days after the last extinction session, sustained threat memory was measured with spontaneous recovery, renewal, and reinstatement. All three tests consisted of 4 CS presentations (60 s inter-trial interval) after a 180 s habituation period. Spontaneous recovery was assayed in the extinction context. Two hours later, renewal was assayed in the conditioning context. Two hours later, mice received two reminder shocks (0.5 mA, 2 s) in the conditioning context, and reinstatement was assayed 24 hrs later in the conditioning context. Extinction blocks, spontaneous recovery, renewal, and reinstatement data are presented as average freezing across four consecutive CSs.

Stimulus delivery and automated analysis of freezing were conducted with Actimetrics FreezeFrame V4 software (Coulbourn). Individual thresholds for freezing were manually set for each animal and behavioral test by determining the highest movement index value that showed no movement by the mouse except breathing for ≥ 1 s. Freezing during threat extinction and recall tests was binned into four CS trial blocks, and pre-CS baseline freezing was analyzed in a separate statistical model. Although freezing is the conventional measure of threat in rodents (Blanchard and Blanchard, 1969; Fanselow, 1980), published studies have demonstrated both strain and sex differences in passive (i.e. freezing) versus active (i.e. flight, grooming) threat responses (Griebel et al., 1997; Gruene et al., 2015a; Tipps et al., 2014). Therefore, CS-evoked flight and grooming behaviors during extinction were scored manually. The flight score was calculated as the total number of rapid darts across the operant chamber, swift circular spins, and jumps (Mongeau et al., 2003). Grooming was defined as active engagement in elliptical stroke, unilateral stroke, bilateral stroke, and/or body licking (Berridge et al., 2005).

### Statistical Analyses

Statistical analyses were conducted with IBM SPSS (Armonk, NY), and data were graphed with GraphPad Prism (San Diego, CA). Normal distribution and homogeneity of variance were tested before proceeding to the appropriate parametric or non-parametric tests. Two-tailed independent-samples t-tests or Mann-Whitney U tests were used to compare two groups with one independent variable. T-tests that failed Levene’s test of equal variances were performed with Welch’s adjustment. One-way analysis of variance (ANOVA) or Kruskal-Wallis tests were used for one dependent variable and ≥ 3 groups. Two-way repeated-measures ANOVA was used for repeating dependent variables and ≥ 2 groups. For repeated-measures tests that violated assumptions of sphericity, Greenhouse-Geisser values were reported. To determine differences in cumulative probability of CS-evoked freezing, grooming, and flight scores, we collapsed the instance of these behaviors across both extinction sessions and used Kolmogorov-Smirnov (K-S) tests to determine significant differences between group distributions. For K-S tests, Bonferroni-adjusted p-values are reported to correct for multiple testing. Categorical data were analyzed with Χ^2^ test for independence. C57Bl/6J, the most commonly used inbred mouse line, was used as the comparison group for all post-hoc analyses. For all tests, familywise α was maintained at 0.05. Effect sizes are reported as Cramér’s V (φ_c_) for Χ^2^ test for independence, eta-squared (η^2^) for ANOVA and Kruskal-Wallis tests, Cohen’s D (d) for t-tests and Bonferroni-adjusted post-hoc comparisons, and Pearson’s coefficient (r) for Mann-Whitney U and Dunn post-tests. Litter effects were tested with one-way and repeated-measured ANCOVA and were not found to contribute to the group differences reported in this manuscript. Full statistics for all analyses are reported in supplementary tables that accompany each figure (Supplementary Tables 1-7). Given the low sample sizes for some estrous cycle comparisons, we additionally provide 1-β from reverse power analyses for every statistically significant comparison in the Results section.

## RESULTS

### Strain differences in estrous cycle regularity and length

As ovulatory cycles have been found to be genetically encoded (Nelson et al., 1992), estrous cycle length and frequency was evaluated across the four mouse strains. In all strains, some females exhibited irregular estrous cycles, defined as ≥5 days in the same cycle stage. However, fewer C57Bl/6J mice exhibited irregular estrous cycles than the other 3 strains (Fig. 1A; X^2^ test: C57Bl/6J vs 129S1/SvlmJ, X^2^ = 12.62, *p* = 0.002, 1-β = 0.607; C57Bl/6J vs DBA/2J, X^2^ = 4.462, *p* < 0.001, 1-β = 0.560; C57Bl/6J vs BALB/cJ, X^2^ = 11.55, *p* = 0.003;, 1-β = 0.568). Cycle length, defined as the number of days from proestrus to proestrus, was quantified in females that completed ≥2 cycles over the 16 testing days. Cycle length also differed across strains (Kruskall-Wallis: H = 8.502, *p* = 0.036, 1-β = 0.567) with longer cycle length in 129S1/SvlmJ compared to C57Bl/6J mice (Fig. 1B; Dunn’s post-test: *p* = 0.047). For all subsequent behavioral data, females with irregular estrous cycles were included in comparisons to males but excluded in comparisons across the estrous cycle.

**Figure 1.**
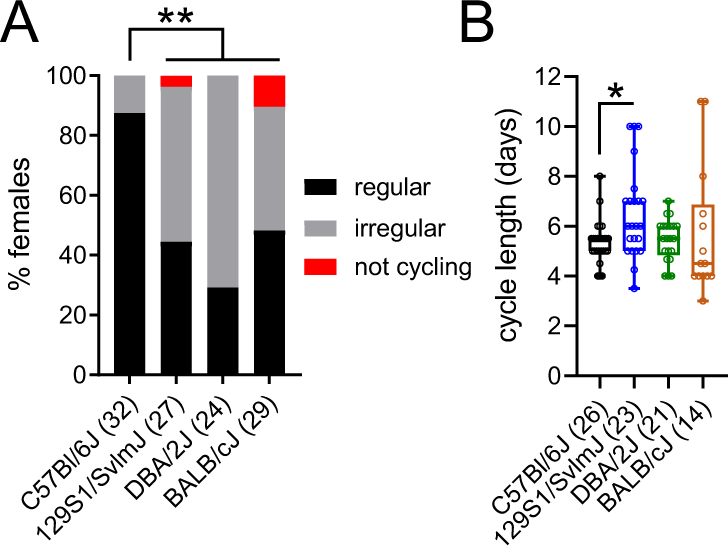
Strain differences in estrous cyclicity. Fewer C57Bl/6J females exhibited irregular estrous cycles than 129S1/SvlmJ, DBA/2J, or BALB/cJ mice (**A**). Cycle length was longer in 129S1/SvlmJ than C57Bl/6J females (**B**). A: χ^2^ test for independence with Bonferroni-corrected *p*-values. B: Kruskal-Wallis test followed by Dunn’s post-test. \**p* < 0.05. n/group denoted in parentheses on X axis labels. Data presented as box-and-whisker plots in B. For statistical details, see Supplementary Table 1.

### Locomotor activity and avoidance behavior varies across mouse strains

To assess responses to potential threat, we quantified locomotor activity and center avoidance in a 30 min open field test and open arm avoidance in a 5 min elevated plus maze test. Significant differences between strains were observed for both measures of the open field test. In both males (Fig. 2A; Kruskall-Wallis: H = 24.00, *p* < 0.001, 1-β = 1.000; Dunn’s post-test: C57Bl/6J vs 129S1/SvlmJ, *p* < 0.001 and C57Bl/6J vs BALB/cJ, *p* < 0.001) and females (Fig. 2F; Kruskall-Wallis: H = 74.12, *p* < 0.001, 1-β = 1.000; Dunn’s post-test: C57Bl/6J vs 129S1/SvlmJ, *p* < 0.001 and C57Bl/6J vs BALB/cJ, *p* < 0.001), 129S1/SvImJ and BABL/cJ mice exhibited reduced locomotor activity compared to C57Bl/6J mice, whereas DBA/2J and C57Bl/6J mice did not differ. Similar strain differences were observed for avoidance behavior, as assayed by the amount of time each mouse spent in the center of the open field. In both males (Fig. 2B-C; Kruskall-Wallis: H = 38.37, *p* < 0.001, 1-β = 1.000; Dunn’s post-test: C57Bl/6J vs 129S1/SvlmJ, *p* < 0.001 and C57Bl/6J vs BALB/cJ, *p* < 0.001) and females (Fig. 2G-H; Kruskall-Wallis: H = 64.09, *p* < 0.001, 1-β = 1.000; Dunn’s post-test: C57Bl/6J vs 129S1/SvlmJ, *p* < 0.001 and C57Bl/6J vs BALB/cJ, *p* < 0.001), 129S1/SvImJ and BALB/cJ mice spent less time in the center of the open field compared to C57Bl/6J mice, whereas DBA/2J and C57Bl/6J mice did not differ. In the elevated plus maze, significant differences between strains were observed for time spent in the open arms in both males (Fig. 2D-E; Kruskall-Wallis: H = 25.46, *p* < 0.001, 1-β = 1.000) and females (Fig. 2I-J; Kruskall-Wallis: H = 22.21, *p* < 0.001, 1-β = 1.000). However, unlike the open field test, post-hoc comparisons revealed that strain differences in this measure of avoidance varied by sex. In males, BALB/cJ (Dunn’s post-test: *p* < 0.001) spent less time in the open arms than C57Bl/6J, whereas 129S1/SvlmJ, DBA/2J, and C57Bl/6J mice did not differ. In females, on the other hand, all strains spent less time in the open arms than C57Bl/6J (Dunn’s post-test: C57Bl/6J vs 129S1/SvlmJ, *p* < 0.001; C57Bl/6J vs DBA/2J, *p* = 0.002; C57Bl/6J vs BALB/cJ, *p* < 0.001). Taken together, these results reveal that strain differences in avoidance behavior elicited by potential threat vary by assay as well as sex.

**Figure 2.**
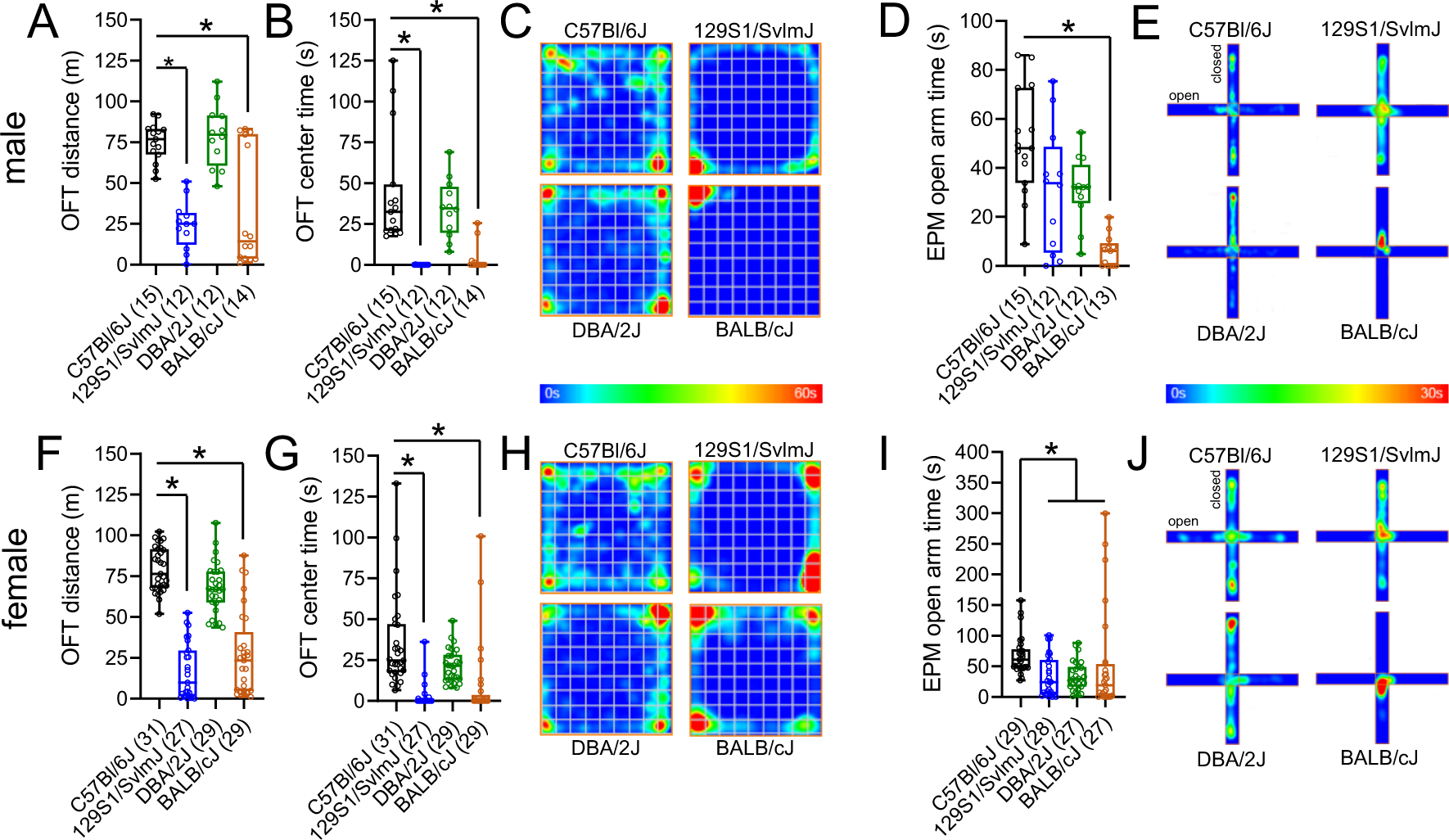
Strain differences in locomotor activity and avoidance. In the open field test (OFT), 129S1/SvlmJ and BALB/cJ mice exhibited reduced locomotor activity and center time compared to C57Bl/6J in both males (**A-B**, representative heat maps **C**) and females (**F-G**, representative heat maps **H**). In the elevated plus maze (EPM) test, male C57Bl/6J mice spent more time in the open arms than BALB/cJ (**D**, representative heat maps **E**), whereas female C57Bl/6J mice spent more time in the open arms than all other strains (**I**, representative heat maps **J**). Kruskal-Wallis test followed by Dunn’s post-test, \**p* < 0.05 as compared with C57Bl/6J. n/group denoted in parentheses on X axis labels. Data presented as box-and-whisker plots. For statistical details, see Supplementary Table 2.

### Fluctuations in avoidance behavior across the estrous cycle are strain-dependent

As cycling ovarian hormones are known to influence avoidance behavior (Maeng and Milad, 2015), we next stratified open field and elevated plus maze data by estrous cycle to make direct within-strain comparisons. In C57Bl/6J mice, open field locomotor activity did not vary between sexes or across the estrous cycle (Fig. 3A), and no differences between males and females were found for time spent in the center of the open field (Fig. 3B). However, C57Bl/6J females in proestrus spent more time in the center of the open field than those in diestrus (Kruskall-Walllis: H = 10.550, *p =* 0.005, 1-β = 0.918; Dunn’s post-test: *p* = 0.001), with a trending difference between proestrus and estrus (Fig. 3B-C; Dunn’s post-test: *p* = 0.051). In the elevated plus maze, females spent more time on the open arms than males (Fig. 3D-E; Welch’s unpaired t-test: t _(37.63)_ = 2.152, *p* = 0.038, 1-β = 0.481). This effect was largely driven by females in proestrus (Kruskall-Wallis: H = 6.800, *p =* 0.033, 1-β = 0.561), which spent more time on the open arms than those in estrus (Dunn’s post-test: *p* = 0.030) or diestrus (Dunn’s post-test: *p* = 0.037).

**Figure 3.**
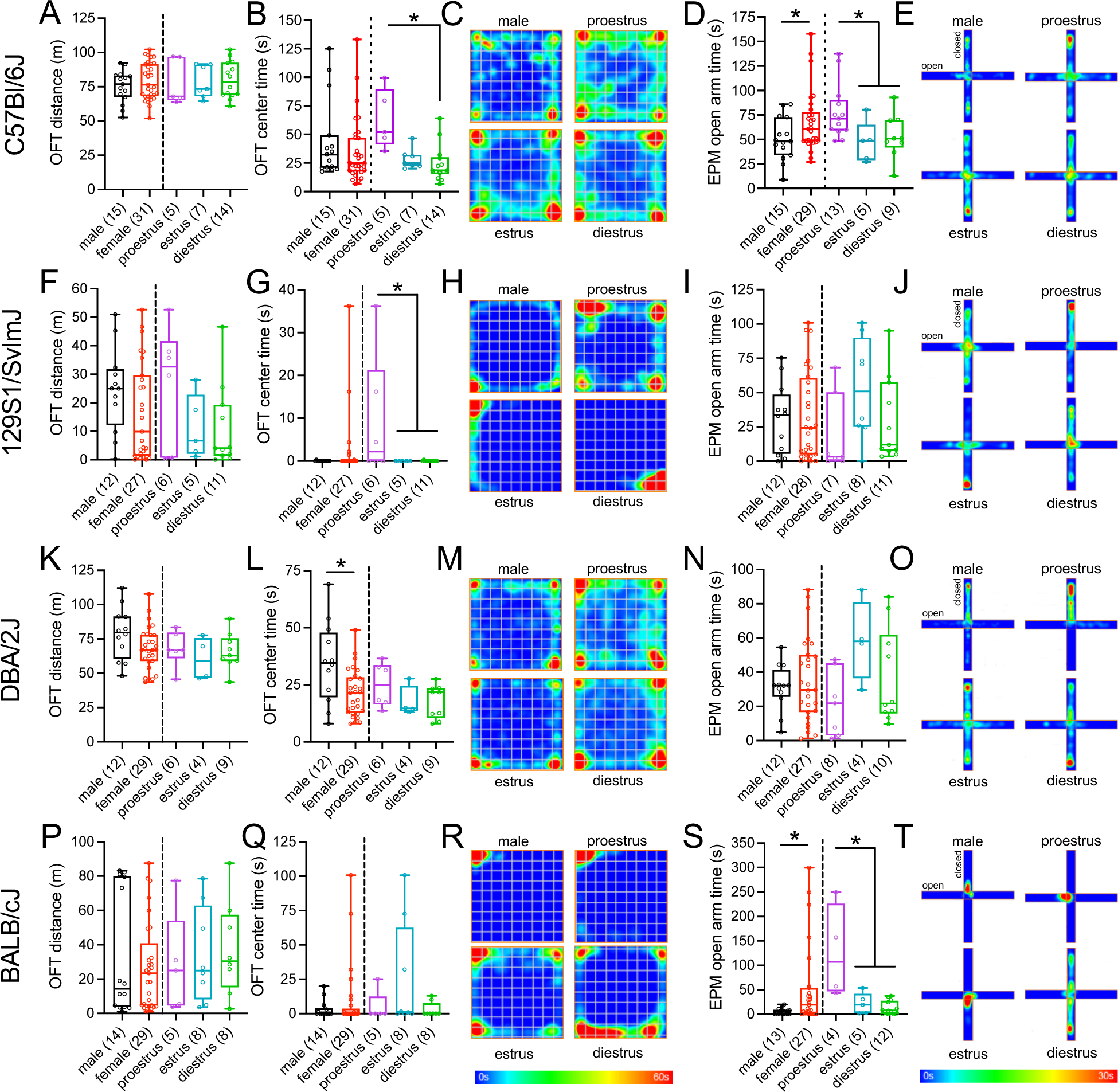
Strain differences in estrous cycle regulation of avoidance behavior. In C57Bl/6J mice, locomotor activity in the open field test (OFT) did not differ between sexes or across the estrous cycle (**A**), but females in proestrus spent more time in the center of the open field than females in diestrus (**B**, representative heat maps **C**). Females spent more time in the open arms of the elevated plus maze (EPM) than males, an effect largely driven by females in proestrus, which spent more time on the open arms than females in estrus or diestrus (**D**, representative heat maps **E**). In 129S1/SvlmJ mice, no differences were observed for OFT locomotor activity (**F**), but females in proestrus spent more time in the center than those in estrus or diestrus (**G**, representative heat maps **H**). No differences were observed for EPM open arm time (**I**, representative heat maps **J**). In DBA/2J mice, no differences were observed for OFT locomotor activity (**K**), but females spent less time in the center than males, independent of estrous cycle stage (**L**, representative heat maps **M**). No differences were observed for EPM open arm time (**N**, representative heat maps **O**). In BALB/cJ mice, no differences were observed in OFT distance or center time (**P-Q**, representative heat maps **R**). Females spent more time on the open arms of the EPM than males, and females in proestrus spent more time on the open arms than females in estrus or diestrus (**S**, representative heat maps **T**). Two-tailed independent-samples t-test/Mann-Whitney U test (male versus female) or one-way ANOVA/Kruskal-Wallis test followed by Dunn’s post-test (estrous cycle comparisons), \**p* < 0.05. n/group denoted in parentheses on X axis labels. Data presented as box-and-whisker plots. For statistical details, see Supplementary Table 3.

In 129S1/SvImJ mice, open field locomotor activity did not vary between sexes or across the estrous cycle (Fig. 3F), and no differences between males and females were found for time spent in the center of the open field (Fig. 3G-H). However, 129S1/SVlmJ females in proestrus spent more time in the center of the open field (Kruskall-Wallis: H = 6.210, *p =* 0.040, 1-β = 0.533) than those in estrus (Dunn’s post-test: *p* = 0.030) or diestrus (Dunn’s post-test: *p* = 0.028). Reduced avoidance in proestrus did not extend to the elevated plus maze, where no differences between sexes or across the estrous cycle were observed for time spent on the open arms (Fig. 3I-J).

In DBA/2J mice, open field locomotor activity did not vary between sexes or across the estrous cycle (Fig. 3K). However, females spent less time in the center of the open field than males (Welch’s unpaired t-test: t _(14.16)_ = 2.403, *p* = 0.031, 1-β = 0.829), an effect not dependent on the estrous cycle (Fig. 3L-M). No difference between sexes or across the estrous cycle were observed for open arm time on the elevated plus maze (Fig. 3N-O).

In BALB/cJ mice, open field locomotor activity and center time did not differ between sexes or across the estrous cycle (Fig. 3P-R). However, females spent more time on the open arms of the elevated plus maze than males (Fig. 3S-T; Mann Whitney: U = 202, *p* = 0.990, 1-β = 0.170). This effect was driven by females in proestrus (Kruskall-Wallis: H = 9.26, *p* = 0.004, 1-β = 0.881), which spent more time on the open arms than those in estrus (Dunn’s post-test: *p* = 0.040) or diestrus (Dunn’s post-test: *p* = 0.002).

These results are in alignment with previous studies in C57Bl/6J mice demonstrating reduced avoidance in the proestrus phase of the estrus cycle, when estrogen, progesterone, follicle stimulating hormone, and luteinizing hormone rapidly surge and peak (Francois et al., 2022; Frye and Rhodes, 2008; Jaric et al., 2019a; Jaric et al., 2019b; Koonce et al., 2012; Walf et al., 2009). However, our data suggest that this effect of proestrus is both strain- and assay-dependent: C57Bl/6J females exhibited a cross-assay decrease in avoidance behavior in proestrus, whereas this effect was only present in the open field test for 129S1/SVlmJ and the elevated plus maze for BALB/cJ. Interestingly, estrous cycle did not impact avoidance behavior in DBA/2J females. Taken together, these results indicate that cycling sex hormones may interact with genetic complement to guide avoidance behavior.

### Conditioned threat memory dynamics vary across mouse strains

We next assayed behavioral responses to acute and sustained threat with auditory threat conditioning and extinction training across the four inbred strains. As DBA/2J and BALB/cJ mice are reported to have very low conditioned threat responses (Balogh and Wehner, 2003; Hefner et al., 2008; Stiedl et al., 1999), we implemented a high-threshold training paradigm consisting of six CS-US pairings in order to make direct comparisons across all four strains. Mice then underwent two days of massed extinction training in an altered context to attenuate the CS-evoked threat memory. We assayed sustained threat memory seven days after the last extinction session with spontaneous recovery in the extinction context, renewal in the conditioning context, and reinstatement in the conditioning context after re-exposure to the US. We found that strain differences in both acute and sustained threat memory vary between the sexes.

In males, DBA/2J and BALB/cJ mice exhibited lower conditioned freezing than C57Bl/6J mice throughout the threat conditioning and extinction training paradigms, whereas 129S1/SVlmJ mice were indistinguishable from C57Bl/6J (Fig. 4A). During threat conditioning (repeated-measures ANOVA: main effect of strain, F_(3,66)_ = 27.95, *p* < 0.001, 1-β = 1.000), DBA/2J and BALB/cJ mice acquired conditioned freezing to the CS less rapidly than C57Bl/6J mice (Bonferroni post-tests: *p* < 0.001). We assayed acute threat memory recall in an altered context the following day. DBA/2J and BALB/cJ mice exhibited lower pre-CS baseline freezing than C57Bl/6J mice (one-way ANOVA: F_(3,66)_ = 17.733, 1-β = 1.000; Bonferroni post-tests: *p* < 0.001). This effect persisted throughout the first day of extinction training (repeated-measures ANOVA: main effect of strain, F_(3,66)_ = 70.24, *p* < 0.001, 1-β = 1.000; Bonferroni post-tests: *p* < 0.001). Correspondingly, on the second day of extinction training, DBA/2J and BALB/cJ mice again exhibited lower pre-CS baseline freezing (one-way ANOVA, F_(3,66)_ = 23.547, *p* < 0.001, 1-β = 1.0; Bonferroni post-tests: *p* < 0.001) and cued-evoked freezing throughout all extinction blocks (repeated-measures ANOVA: main effect of strain, F_(3,66)_ = 67.270, *p* < 0.001, 1-β = 1.000; Bonferroni post-tests: *p* < 0.001) compared to C57Bl/6J mice. Seven days later, we measured sustained threat memory and found that DBA/2J and BALB/cJ mice exhibited lower spontaneous recovery (one-way ANOVA: F_(3,66)_ = 27.725, *p* < 0.001, 1-β = 1.000; Bonferroni post-tests: *p* < 0.001) and renewal (one-way ANOVA: F_(3,66)_ = 33.351, *p* < 0.001, 1-β = 1.000; Bonferroni post-tests: *p* < 0.001) than C57Bl/6J mice. However, whereas DBA/2J mice did not exhibit threat memory reinstatement 24 hrs following reminder shocks (one-way ANOVA: F_(3,66)_ = 30.975, *p* < 0.001, 1-β = 1.000; Bonferroni post-tests: *p* < 0.001), BALB/cJ males exhibited memory reinstatement that was indistinguishable from C57Bl/6J.

**Figure 4.**
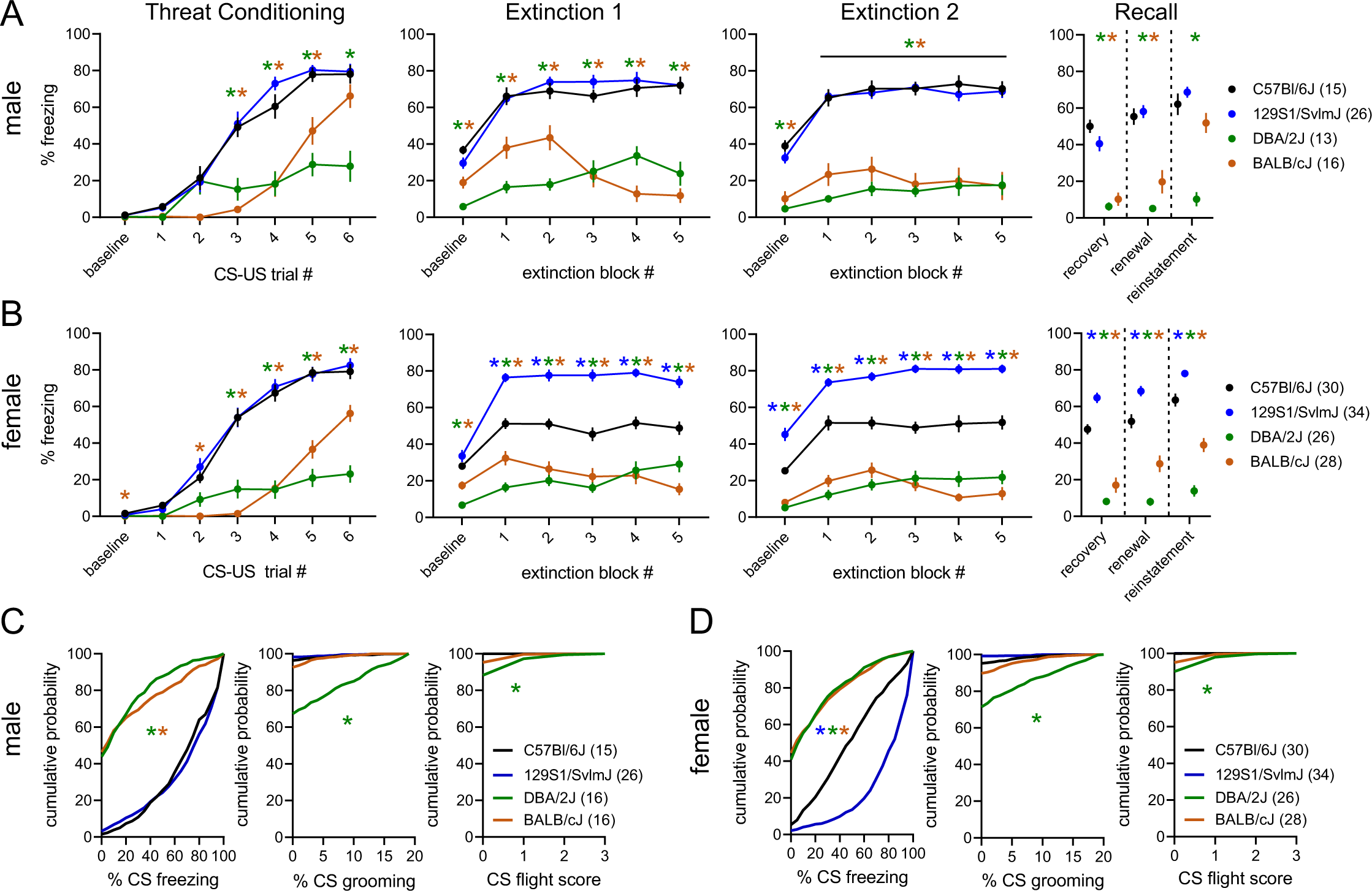
Strain differences in auditory threat memory dynamics. In males, DBA/2J and BALB/cJ acquired CS-evoked freezing less rapidly than C57Bl/6J during auditory threat conditioning. Reduced freezing in DBA/2J and BALB/cJ compared to C57Bl/6J continued during the pre-CS baseline period and CS blocks of the first and second extinction sessions, as well as during spontaneous recovery and renewal. Only DBA/2J froze less than C57Bl/6J during reinstatement (**A**). In females, DBA/2J and BALB/cJ mice acquired cue-evoked freezing less rapidly than C57Bl/6J during auditory threat conditioning. Compared to C57Bl/6J, DBA/2J and BALB/cJ froze less, whereas 129S1/SvlmJ froze more, during the pre-CS baseline periods and all CS blocks of the first and second extinction sessions, as well as during spontaneous recovery, renewal, and reinstatement (**B**). Quantification of CS-evoked behavior collapsed across extinction sessions (**C-D**). Freezing and grooming data are presented as percent time during CS presentations, and the flight score is presented as the total number of CS-evoked darts, spins, and jumps. In males, DBA/2J and BALB/cJ froze less than C57Bl/6J, and DBA/2J spent more time engaged in grooming and flight behavior during CS presentations (**C**). In females, DBA/2J and BALB/cJ females froze less and 129S1/SvlmJ froze more than C57Bl/6J. DBA/2J females spent more time engaged in grooming and flight behavior during CS presentations (**D**). A-B: two-way repeated-measures ANOVA followed by Bonferroni-corrected post-hoc comparisons in the case of a significant interaction (threat conditioning, extinction blocks) or one-way ANOVA followed by Bonferroni-corrected post-hoc comparisons (baseline, recovery, renewal, reinstatement). C-D: Kolmogorov–Smirnov tests with Bonferroni-corrected *p* values. \**p* < 0.05 for C57Bl/6J versus 129S1/SvlmJ (blue asterisks), DBA2/J (green asterisks), and BALB/cJ (gray asterisks). n/group denoted in parentheses on legends. Data presented as mean ± SEM (A-B) or cumulative probability plots (C-D). For statistical details, see Supplementary Table 4.

In females, on the other hand, threat memory dynamics differed between C57Bl/6J mice and the three other strains (Fig. 4B). During threat conditioning (repeated-measures ANOVA: main effect of strain, F_(3,114)_ = 69.517, *p* < 0.001, 1-β = 1.000), DBA/2J and BALB/cJ mice acquired conditioned freezing to the CS less rapidly than C57Bl/6J mice (Bonferroni post-tests: *p* < 0.001). Acute threat memory recall was assayed in a novel context the following day. DBA/2J and BALB/cJ froze less than C57Bl/6J during the pre-CS baseline (one-way ANOVA: F_(3,114)_ = 23.562, *p* < 0.001, 1-β = 1.000, Bonferroni post-tests: *p* < 0.01). Throughout the first day of extinction training (repeated-measures ANOVA: main effect of strain, F_(3,114)_ = 108.691, *p* < 0.001, 1-β = 1.000), 129S1/SvImJ froze more than, and DBA/2J and BALB/cJ less than, C57Bl/6J during CS presentations (Bonferroni post-tests: *p* < 0.001). Correspondingly, on the second day of extinction training, DBA/2J and BALB/cJ froze less than, and 129S1/SvImJ froze more than, C57Bl/6J mice during the pre-CS baseline period (one-way ANOVA: F_(3,114)_ = 57.099, *p* < 0.001, 1-β = 1.000; Bonferroni post-tests: *p* < 0.001) and to the CS throughout the five extinction blocks repeated-measures ANOVA: main effect of strain, F_(3,114)_ = 137.801, *p* < 0.001, 1-β = 1.000; Bonferroni post-tests: *p* < 0.001). These strain-dependent effects persisted when sustained threat memory was assayed one week later. DBA/2J and BALB/cJ exhibited less, and 129S1/SvImJ more, cue-evoked freezing than C57Bl/6J females during spontaneous recovery (one-way ANOVA: F_(3,114)_ = 77.465, *p* < 0.001, 1-β = 1.000; Bonferroni post-tests: *p* < 0.001), renewal (one-way ANOVA: F_(3,114)_ = 59.529, *p* < 0.001, 1-β = 1.000; Bonferroni post-tests: *p* < 0.002), and reinstatement (one-way ANOVA: F_(3,114)_ = 69.529, *p* < 0.001, 1-β = 1.000; Bonferroni post-tests: *p* < 0.002).

Given the robust differences between strains on post-conditioning cue-evoked freezing, we questioned whether strains with low freezing have a genetic propensity toward active (i.e. flight) rather than passive (i.e. freezing) conditioned defensive responses. We therefore manually quantified a repertoire of behaviors observed during CS presentations throughout both extinction sessions and found that mice typically engaged in three conditioned behavioral responses: freezing, grooming, and flight. We collapsed instances of these behaviors across all 40 CSs of both extinction sessions and quantified their cumulative probability across mouse strains. Mirroring our previous results, strain differences in the cumulative distribution of cue-evoked freezing were sex-dependent. In males DBA/2J (K-S test: D = 0.659, *p* < 0.001) and BALB/cJ (K-S test: D = 0.591, *p* < 0.001), but not 129S1/SvImJ, differed from C57Bl/6J (Fig. 4C), whereas in females all three strains differed from C57Bl/6J (Fig. 4D; K-S test: D > 0.445, *p* < 0.001). Consistent with low freezing throughout extinction sessions as well as prior publications (Griebel et al., 1997; Tipps et al., 2014), DBA/2J males and females engaged in more grooming (K-S test: D > 0.230, *p* < 0.001) and flight (K-S test: D > 0.098, *p* < 0.001) during CS presentations than sex-matched C57Bl/6J mice. These data suggest that both acute and sustained threat memory, as well as the modality of cue-evoked defensive behavioral responses, are genetically encoded.

### Sex differences in threat memory dynamics are strain-dependent

Finally, we stratified our threat conditioning data by sex and estrous cycle to make direct within-strain comparisons. In C57Bl/6J mice, males and females acquired conditioned freezing at the same rate (Fig. 5A), in accordance with a previous report implementing this high-threshold paradigm in this mouse line (du Plessis et al., 2022). When acute threat memory recall was assessed in an altered context the following day, females froze less than males during both the pre-CS baseline (unpaired t-test: t_(43)_ = 3.041, *p* = 0.004, 1-β = 0.860) and CS presentations throughout the entire first session of extinction (repeated-measures ANOVA: main effect of sex, F_(1,43)_ = 21.695, *p* < 0.001, 1-β = 0.996). The following day, females continued to freeze less than males during the pre-CS baseline (unpaired t-test: t_(43)_ = 3.875, *p* < 0.001, 1-β = 0.972) and CS presentations throughout the second session of extinction (repeated-measures ANOVA: main effect of sex, F_(1,43)_ = 15.451, *p* < 0.001, 1-β = 0.975). However, no sex differences were observed on sustained threat memory when spontaneous recovery, renewal, and reinstatement were assessed one week later. These sex differences in threat memory dynamics were not driven by estrous cycle stage during threat conditioning (Supplementary Fig. 1A) or the first extinction session (Supplementary Fig. 2A). However, females conditioned in estrous exhibited reduced spontaneous recovery compared to those conditioned in proestrus (one-way ANOVA: F_(2,26)_ = 4.098, *p* = 0.028, 1-β = 0.726; LSD post-test: *p* = 0.011) or diestrus (Supplementary Fig. 1A; LSD post-test: *p* = 0.034). We questioned whether sex differences in active versus passive conditioned responses could account for reduced freezing in females. However, when evaluating behavior across CS presentations of both extinction sessions, sex differences in freezing (K-S test: D = 0.295, *p <* 0.001) but not grooming or flight were observed (Fig. 5B). These data suggest that C57Bl/6J males initially encode stronger threat memories than females but that sustained threat memory is similar between the sexes.

**Figure 5.**
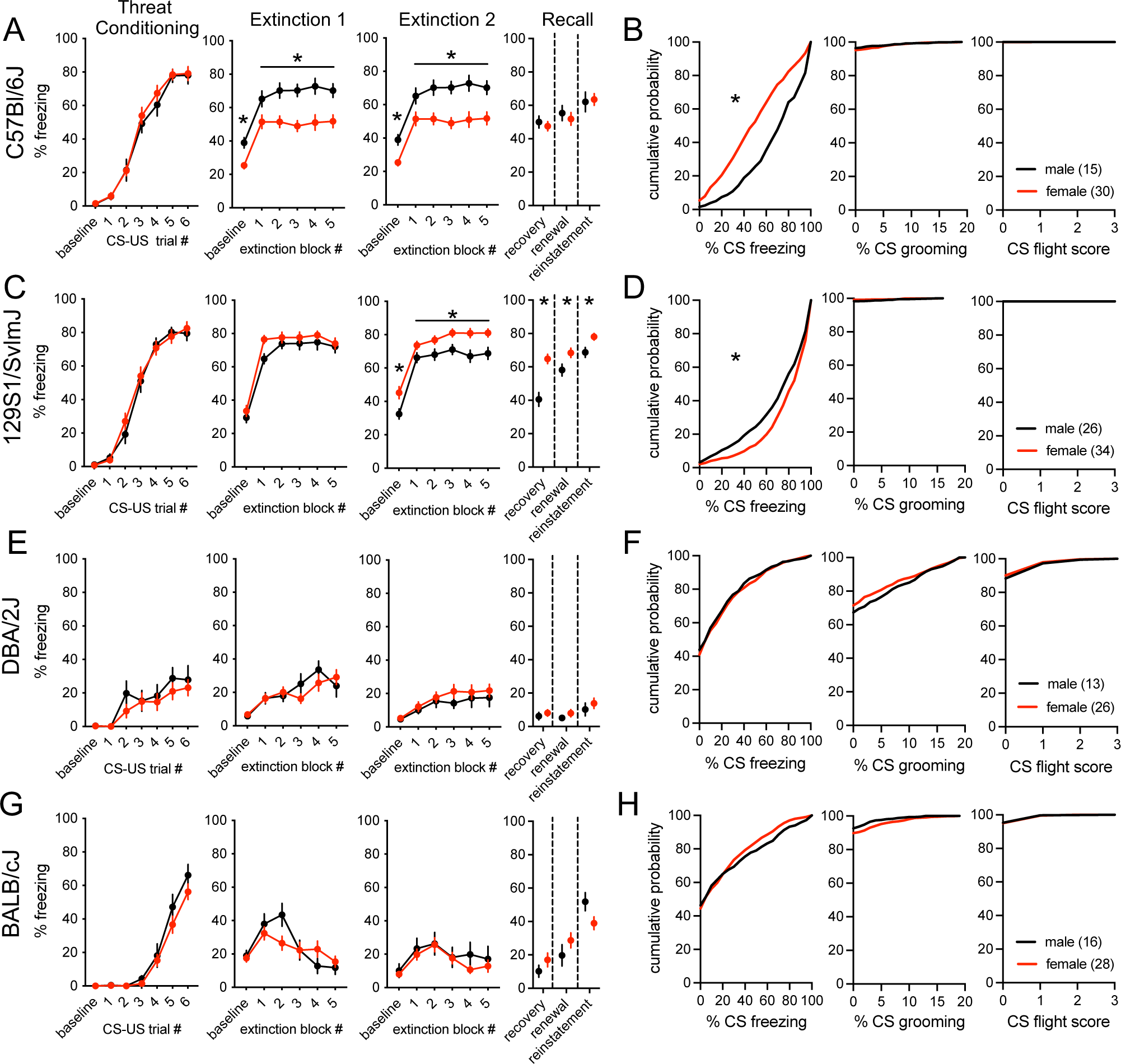
Sex differences in auditory threat memory dynamics are strain-dependent. In C57Bl/6J, male and female mice acquired CS-evoked freezing at the same rate during threat conditioning, but females froze less than males during the pre-CS baseline period and CS blocks of the first and second extinction sessions. No sex differences were observed in spontaneous recovery, renewal, or reinstatement of threat memory (**A**). Quantification of CS-evoked behavior collapsed across extinction sessions revealed that C57Bl/6J females froze less than males, while limited grooming or flight behavior was observed in either sex (**B**). In 129S1/SvlmJ, male and female mice acquired CS-evoked freezing at the same rate during threat conditioning and exhibited similar baseline and CS-evoked freezing during the first extinction session. However, females exhibited greater sustained threat memory, freezing more than males during the pre-CS baseline period and CS-blocks of the second extinction session as well as during spontaneous recovery, renewal, and reinstatement (**C**). Quantification of cue-evoked behavior collapsed across both extinction sessions revealed that 129S1/SvlmJ females froze more than males, while limited grooming or flight behavior was observed in either sex (**D**). In DBA/2J, no sex differences in auditory threat memory dynamics were observed (**E**). CS-evoked behavioral responses across extinction sessions did not differ between male and female DBA/2J mice (**F**). In BALB/cJ, no sex differences in auditory threat memory dynamics were observed (**G**). CS-evoked behavioral responses across extinction sessions did not differ between male and female BALB/cJ mice (**H**). A,C,E,G: two-way repeated-measures ANOVA followed by Bonferroni-corrected post-hoc comparisons in the case of a significant interaction (threat conditioning, extinction blocks) or two-tailed independent-samples t-tests (baseline, recovery, renewal, reinstatement). B,D,F,H: Kolmogorov–Smirnov tests. \**p* < 0.05. n/group denoted in parentheses in legends. Data presented as mean ± SEM (A,C,E,G) or cumulative probability plots (B,D,F,H). For statistical details, see Supplementary Table 5.

In 129S1/SvImJ mice, males and females also acquired conditioned threat at the same rate (Fig. 5C). Males and females were again indistinguishable during pre-CS baseline and CS-evoked freezing during the first extinction session. However, sex differences emerged during the second extinction session with females freezing more during the pre-CS baseline period (Welch’s t-test: t_(57.94)_ = 2.695, *p* = 0.012, 1-β = 0.737) and throughout blocks of CS presentations (repeated-measures ANOVA: main effect of sex, F_(1,58)_ = 18.399, *p* < 0.001, 1-β = 0.990). This phenotype resembled an incubation of threat memory responses in females (Eysenck, 1968), evidenced by enhanced spontaneous recovery (Welch’s t-test: t_(45.53)_ = 4.858, *p* < 0.001, 1-β = 0.999), renewal (unpaired t-test: t_(58)_ = 2.334, *p* = 0.023, 1-β = 0.647), and reinstatement (unpaired t-test: t_(58)_ = 2.700, *p* = 0.009, 1-β = 0.772). Similar to C57Bl/6J mice, estrous cycle stage during threat conditioning (Supplementary Fig. 1B) or the first extinction session (Supplementary Fig. 2B) did not account for sex differences in threat memory dynamics. However, females undergoing threat conditioning or the first day of extinction training in diestrus exhibited enhanced spontaneous recovery (one-way ANOVA: F_(2,26)_ = 4.586, *p* = 0.020, 1-β = 0.776; LSD post-test: vs proestrus, *p* = 0.016, vs estrus, *p =* 0.032). Sex differences in freezing (K-S test: D = 0.124, *p* < 0.001), but not grooming or flight, were observed when CS-evoked behaviors were collapsed across the 40 CSs of both extinction sessions (Fig. 5D). These data suggest that acute threat memory is similar between the sexes, while sustained threat memory is stronger in 129S1/SvlmJ females than males.

In DBA/2J mice, no sex differences (Fig. 5E-F) or estrous-cycle dependent effects (Supplementary Figs. 1C and 2C) were observed in auditory threat memory dynamics.

In BALB/cJ mice, no sex differences were observed in auditory threat memory dynamics, including presentation of passive versus active threat responses (Fig. 5G-H). However, when we stratified data by estrous cycle stage during threat conditioning, we found that BALB/cJ females in estrus formed threat memories that were resistant to extinction (Supplementary Fig. 1D). No estrous cycle dependent effects were observed during threat conditioning or extinction session one, but females conditioned in estrus froze more during baseline (one-way ANOVA: F_(2,17)_ = 4.488, *p* = 0.027, 1-β = 0.762; LSD post-test: *p =* 0.014), throughout the second session of extinction (repeated measures ANOVA, main effect of estrous cycle: F_(2,17)_ = 5.115, *p* = 0.018, 1-β = 0.817; LSD post-test: *p =* 0.012), and exhibited higher reinstatement (one-way ANOVA: F_(2,17)_ = 21.837, *p* < 0.001, 1-β = 1.000; LSD post-test: *p* < 0.001), but not spontaneous recovery or renewal, than those conditioned in diestrus. On the other hand, mice undergoing the first day of extinction learning in diestrus exhibited enhanced reinstatement compared to those in proestrus (one-way ANOVA: F_(2,17)_ = 3.750, *p* = 0.045, 1-β = 0.679; LSD post-test: *p* = 0.018).

Together, these data demonstrate that basal sex differences as well as potential estrous cycle regulation of threat memory dynamics are strain dependent. In C57Bl/6J mice, females exhibit higher acute, but not sustained, threat memory responses compared to males. On the other hand, 129S1/SvImJ females exhibit higher sustained, but not acute, threat responses compared to males. No consistent trends in estrous cycle regulation of threat memory dynamics were observed across strains. Conditioning during estrous decreased spontaneous recovery in the C57Bl/6J and 129S1/SvlmJ lines but impaired extinction learning and enhanced reinstatement in the BALB/cJ line. Contrary to previous findings in rats (Blume et al., 2017; Blume et al., 2019; Bouchet et al., 2017; Chang et al., 2009; Graham and Daher, 2016; Graham and Milad, 2013; Graham and Scott, 2018; Gruene et al., 2015b; Milad et al., 2009; Rey et al., 2014; Voulo and Parsons, 2019; Zeidan et al., 2011), we did not observe enhanced extinction memory consolidation in mice undergoing the first session of extinction training in proestrus in any mouse strain. However, as we did not target behavior to specific estrous stages in these experiments, the proestrus groups in our analysis of threat conditioning dynamics stratified by cycle stage during the first extinction session are underpowered.

## DISCUSSION

By comparing four common inbred mouse strains, we determined that genetic complement contributes to sex differences and estrous cycle fluctuations in behavioral responses related to three components of negative valence systems: avoidance of potential threat in open spaces, acute threat memory prior to extinction training, and sustained threat memory after extinction training. We observed a large genetic contribution to all behaviors under investigation, as well as a surprising genetic dissociation between avoidance and threat memory. Basal sex differences and estrous cycle regulation of avoidance behavior were strain- and assay-dependent. Similarly, basal sex differences in acute and sustained threat memory were strain-dependent, but with limited impact of estrous cycle stage. We expand upon these points in the discussion below and conclude with future directions to further this line of inquiry.

Independent of sex, we found that C57Bl/6J mice exhibited low avoidance but high threat memory, 129S1/SvlmJ mice high avoidance and high threat memory, DBA/2J mice low avoidance and low threat memory, and BALB/cJ mice high avoidance but low threat memory. These findings are largely in accordance with previous studies (Crawley et al., 1997), the majority of which excluded female subjects. Notably, we used a high threshold threat conditioning paradigm to compare acute and sustained threat memory across the four strains, as DBA/2J and BALB/cJ mice exhibit floor effects in conditioned threat responses (Balogh and Wehner, 2003; Hefner et al., 2008; Stiedl et al., 1999). However, this paradigm led to extinction-resistant threat memories in C57Bl/6J mice, which generally exhibit high acute but low sustained threat memory (Camp et al., 2009; Camp et al., 2012; Hefner et al., 2008; Lucas et al., 2019). As previously reported (Clark et al., 2019), this high intensity paradigm also unmasked sex differences in C57Bl/6J mice, whereby females exhibit reduced acute, but equivalent sustained, conditioned threat responses compared to males. We also find that previous reports of high acute and sustained threat memory in 129S1/SvlmJ males are more pronounced in females (Camp et al., 2009; Camp et al., 2012; Hefner et al., 2008). The robust and opposing sex differences between C57Bl/6J and 129S1/SvlmJ mice may be useful in the study of mechanisms driving susceptibility versus resilience to threat memory endophenotypes. Unlike previous reports in rats (Gruene et al., 2015a; Milad et al., 2009), these sex differences were not attributable to estrous cycle phase or divergence of passive versus active defensive behavioral responses, which were not observed in any mouse strain.

Reproductive hormone fluctuations have been shown to influence behavioral responses to potential threat across species (Maeng and Milad, 2015). Across mouse strains, decreased avoidance behavior in the proestrus phase has been reported in some (Datta et al., 2016; Datta et al., 2019; Francois et al., 2022; Galeeva and Tuohimaa, 2001; Jaric et al., 2019a; Jaric et al., 2019b; Koonce et al., 2012; Meziane et al., 2007; Naule et al., 2015; Paris et al., 2014; Walf et al., 2009) but not all (Bath et al., 2012; Kastenberger et al., 2012; Kastenberger and Schwarzer, 2014; Laham et al., 2022; Levy et al., 2023; Manzano-Nieves et al., 2018; Yohn et al., 2020) previous studies. To our knowledge, no publications to date have investigated estrous cycle regulation of avoidance behavior in 129S1/SvlmJ or DBA/2J mice. Here, we demonstrate reduced avoidance behavior in proestrus in C57Bl/6J, 129S1/SvlmJ, and BALB/cJ mice, with C57Bl/6J exhibiting decreased avoidance in both the elevated plus maze and the open field test. As previously reviewed (Kokras and Dalla, 2014), we find that females exhibit less avoidance behavior than males. However, this effect is only observed in C57Bl/6J and BALB/cJ mice in the elevated plus maze, but not open field, and was largely driven by females in proestrus. Interestingly, avoidance behavior was unaffected by the reproductive cycle in DBA/2J mice, although a basal sex difference was observed in the open field with increased avoidance in females compared to males. Our data suggest that the C57Bl/6J line is ideal for the study of estrous cycle regulation of avoidance behavior, as they exhibit the most consistent estrous cyclicity as well as cross-assay behavioral effects that persist to old age (Baumgartner et al., 2023). Ultimately, these findings suggest that the impacts of estrous cycle on avoidance behaviors are highly sensitive to both strain and test.

While our findings indicate that genetic complement interacts with cycling reproductive hormones to guide negative valence endophenotypes, the mechanisms driving these effects remain enigmatic. In humans, many polymorphisms within genes regulating ovarian hormone signaling have been identified (for review, see (Maney, 2017), several of which promote susceptibility and resilience to anxiety-related disorders. Polymorphisms in *ESR1,* the gene encoding estrogen receptor α, are associated with increased risk of anxiety disorders and worsened symptom severity in both men and women (Comings et al., 1999; Levey et al., 2020; Prichard et al., 2002; Ryan et al., 2011; Sundermann et al., 2010; Tiemeier et al., 2005), as well as with increased harm avoidance tendencies in women (Gade-Andavolu et al., 2009; Giegling et al., 2009; Miller et al., 2010). The well-studied XbaI polymorphism in *ESR1* has also been shown to modulate the impacts of cycling ovarian hormone levels across the menstrual cycle on emotional processing in women with premenstrual dysphoric disorder (Yen et al., 2018). Beyond changes in sex hormone receptors, polymorphisms in estrogen or progesterone response elements could also influence sex hormone behavioral regulation. For example, single nucleotide polymorphisms in the estrogen response element of the stress-related gene *ADCYAP1R1* are associated with increased risk of post-traumatic stress disorder in women but not men (Ressler et al., 2011). Finally, polymorphisms in *AKR1C1*, which encodes one of the enzymes necessary to convert progesterone to allopregnanolone, are associated with increased risk of panic disorder in women but not men (Quast et al., 2014). While such clinical evidence suggests strong interactive effects between genetic variation and sex hormone signaling in modulating negative valence behaviors in humans, the specific genetic loci driving the behavioral effects in the present study remain unknown. Previous work in C57Bl/6J mice implicates a role for estrogen receptor β (*Esr2*) as one potential candidate driving estrous cycle regulation of avoidance behavior. Germline knockout of *Esr2*, but not *Esr1*, increases avoidance behavior in females, but not males (Krezel et al., 2001). In alignment with our findings, wildtype littermates exhibit reduced avoidance in proestrus, and this effect is eliminated by germline or conditional nervous system deletion of *Esr2* (Naule et al., 2015; Walf et al., 2009). Future studies employing forward genetics approaches utilizing collaborative cross or diversity outbred mice are required to pinpoint sex-specific and hormone-sensitive genetic loci for regulation of negative valence behaviors.

Before closing, we would like to highlight several limitations in the overall design of the present study. First, despite high sample sizes for sex comparisons, some of the estrous cycle comparisons are underpowered due to strain differences in cycle length and regularity (Fig. 1). Following the considerations described in (Goldman et al., 2007), our laboratory now targets experiments to specific estrous cycle phases following estrous tracking for at least two cycles to ensure adequate sample size. We recommend that other labs interested in estrous cycle as an independent variable follow a similar approach. Second, performance issues limit the interpretations that can be made for some aspects of our threat memory dynamics data. Impaired extinction learning in the C57Bl/6J and 129S1/SvlmJ strains confounds the sustained threat memory assays, and absence of passive defensive behavioral responses in DBA2/J confounds the use of freezing as a proxy for memory in this mouse line. Nonetheless, interesting strain and sex differences emerged from this experimental design. For example, despite lack of extinction C57Bl/6J and 129S1/SvlmJ mice, cued threat memory seems to “fade” with time in males but not in females, as evidenced by reduced freezing during spontaneous recovery compared to the last extinction block. While beyond the scope of the present study, we hope that our findings will guide future work exploring the maximal perimeters required for assessment of threat memory dynamics within each strain (i.e., Cazares et al., 2019; Wimer et al., 1968) in order to better assess of the impacts of sex and estrous cycle. Finally, our experimental design focused on the contributions of estrous cycle to endophenotypes relevant to negative valence systems and offers little insight to sex hormone regulation of behavior in males. Male testosterone is regulated on a 24-hr circadian clock with higher levels in the morning than evening and significant variation across housing conditions and inbred mouse strains (Lucas and Eleftheriou, 1980; Sayegh et al., 1990). Future studies should measure and manipulate hormones levels in both sexes to identify convergence and divergence of mechanisms.

In conclusion, our findings demonstrate pronounced impacts of mouse strain on sex differences and estrous cycle regulation of negative valence behaviors. As such, researchers interested in studying these topics should carefully consider these differences when choosing the optimal animal model for their experimental purposes as well as while interpreting experimental outcomes. Additionally, our findings demonstrate the pitfalls of generalizing behavioral results from one mouse strain to another, which likely contributes to the reproducibility crisis in behavioral neuroscience.

## ACKNOWLEDGEMENTS

Many thanks are due to Arabelis Wally and Shannon Norton for animal care and to Alex Yankoglu for assistance with vaginal cytology categorization.

## AUTHOR CONTRIBUTIONS

**Garret Ryherd:** Methodology, Formal Analysis, Investigation, Writing – Reviewing & Editing **Averie L. Bunce:** Formal Analysis, Investigation, Writing – Reviewing & Editing **Haley A. Edwards:** Formal Analysis, Investigation, Writing – Reviewing & Editing **Nina E. Baumgartner:** Formal Analysis, Writing – Reviewing & Editing **Elizabeth K. Lucas:** Conceptualization, Methodology, Formal Analysis, Investigation, Resources, Writing – Original Draft, Writing – Review & Editing, Supervision, Project Administration, Funding Acquisition.

