## Supplementary Figure 1-2 for "Sex differences in avoidance behavior and cued threat memory dynamics in mice: Interactions between estrous cycle and genetic background"

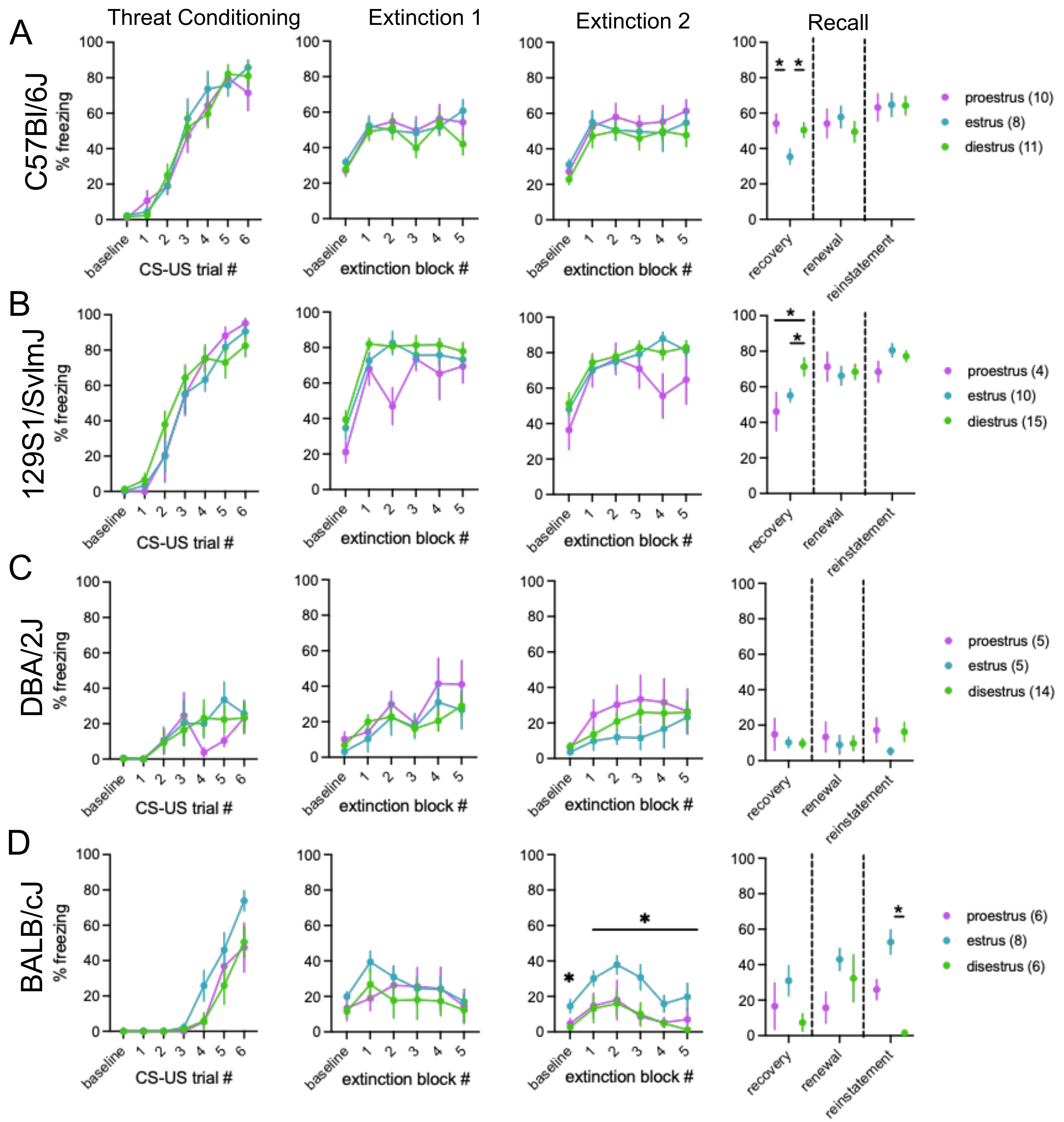


**Supplementary Figure 1.** *Threat memory dynamics stratified by estrous cycle stage during threat conditioning.* In C57Bl/6J, females in estrus during conditioning exhibited reduced spontaneous recovery (**A**). In 129S1/SvlmJ, females in diestrus during conditioning exhibited enhanced spontaneous recovery (**B**). In DBA/2J mice, estrous cycle stage at conditioning did not impact threat conditioning dynamics (**C**). In BALB/cJ, females in estrus during conditioning exhibited increased freezing during the pre-CS baseline period and throughout CS blocks of the second extinction session as well as during threat memory reinstatement (**D**). Two-way repeated-measures ANOVA (threat conditioning, extinction blocks) or one-way ANOVA followed by Fisher's LSD (baseline, recovery, renewal, reinstatement). *$p < 0.05$. n/group denoted in parentheses in legends. Data presented as mean ± SEM. For statistical details, see Supplementary Table 6.

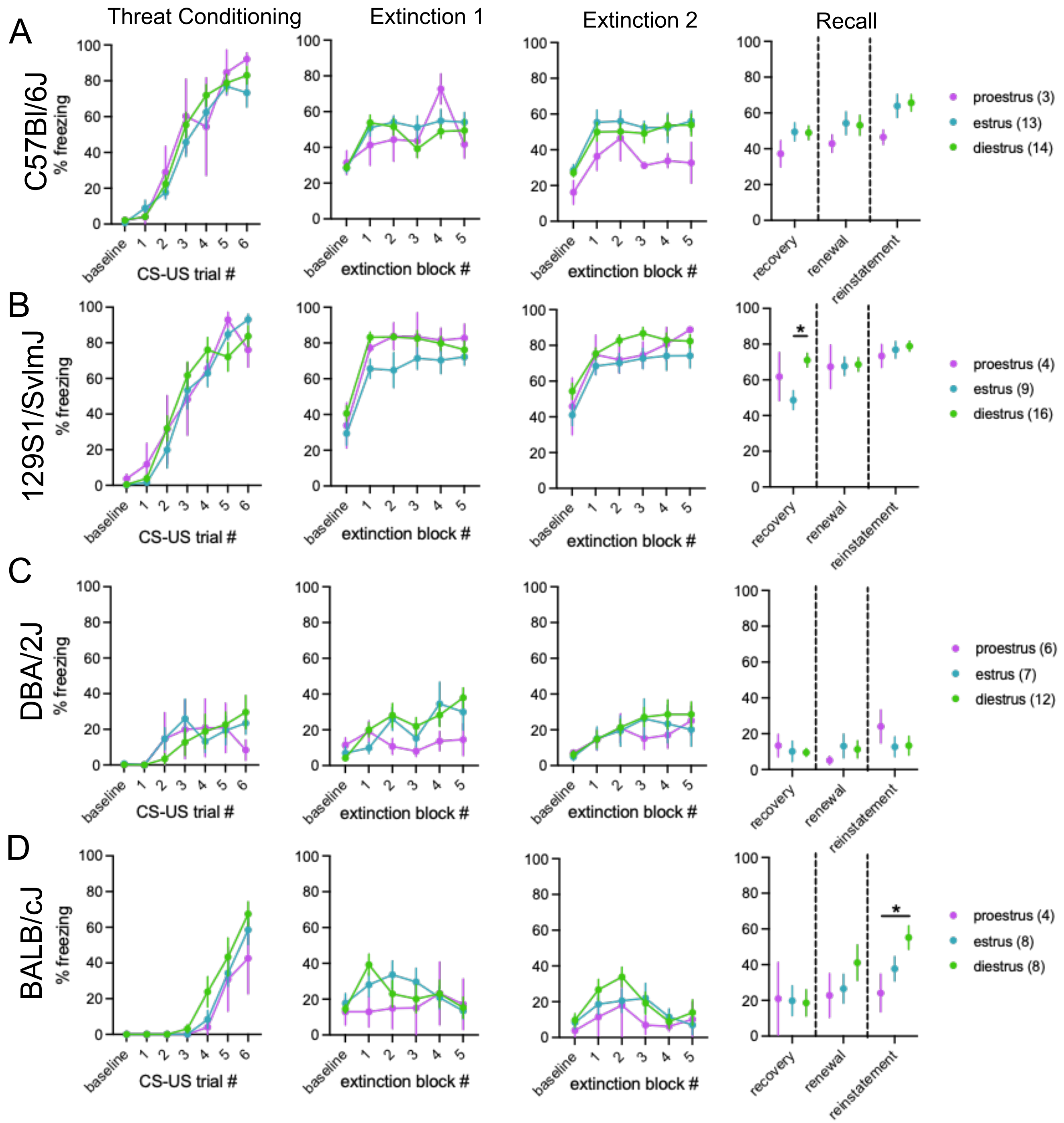


**Supplementary Figure 2.** *Threat memory dynamics stratified by estrous cycle stage during the first extinction session.* In C57Bl/6J, estrous cycle stage during the first extinction session did not impact threat memory dynamics (**A**). In 129S1/SvlmJ, females in estrus during the first extinction session exhibited reduced spontaneous recovery compared to females in diestrus (**B**). In DBA/2J, estrous cycle stage during the first extinction session did not impact threat memory dynamics (**C**). In BALB/cJ, females in proestrus during the first extinction session exhibited reduced reinstatement compared to females in diestrus (**D**). Two-way repeated-measures ANOVA (threat conditioning, extinction blocks) or one-way ANOVA followed by Fisher's LSD (baseline, recall, recovery, renewal, reinstatement). *$p < 0.05$. n/group denoted in parentheses in legends. Data presented as mean ± SEM. For statistical details, see Supplementary Table 7.
